# Improving retinal vascular endothelial cell tropism through rational rAAV capsid design

**DOI:** 10.1101/2023.02.26.530096

**Authors:** Ramesh Periasamy, Dwani D. Patel, Sanford L. Boye, Shannon E. Boye, Daniel M. Lipinski

**Author notes:** Department of Ophthalmology and Visual Science, Medical College of Wisconsin, Milwaukee, Wisconsin, United States. These authors contributed equally to this work (Ramesh Periasamy, Dwani D. Patel). These authors also contributed equally to this work (Sanford L. Boye, Shannon E. Boye, Daniel M. Lipinski).

## Abstract

Vascular endothelial cells (VEC) are essential for retinal homeostasis and their dysfunction underlies pathogenesis in diabetic retinopathy (DR) and exudative age-related macular degeneration (AMD). Studies have shown that recombinant adeno-associated virus (rAAV) vectors are effective at delivering new genetic material to neural and glial cells within the retina, but targeting VECs remains challenging. To overcome this limitation, herein we developed rAAV capsid mutant vectors with improved tropism towards retinal VEC. rAAV2/2, 2/2[QuadYF-TV], and rAAV2/9 serotype vectors (n=9, capsid mutants per serotype) expressing GFP were generated by inserting heptameric peptides (7AA) designed to increase endothelial targeting at positions 588 (2/2 and 2/2[QuadYF-TV] or 589 (2/9) of the virus protein (VP 1-3). The packaging and transduction efficiency of the vectors were assessed in HEK293T and bovine VECs using Fluorescence microscopy and flow cytometry, leading to the identification of one mutant, termed EC5, that showed improved endothelial tropism when inserted into all three capsid serotypes. Intra-ocular and intravenous administration of EC5 mutants in C57Bl/6j mice demonstrated moderately improved transduction of the retinal vasculature, particularly surrounding the optic nerve head, and evidence of sinusoidal endothelial cell transduction in the liver. Most notably, intravenous administration of the rAAV2/2[QuadYF-TV] EC5 mutant led to a dramatic and unexpected increase in cardiac muscle transduction.

## Introduction

Diabetic retinopathy (**DR**) and age-related macular degeneration (**AMD**) are leading causes of blindness worldwide and are characterized by the dysfunction of retinal microvascular endothelial cells (**MVECs**), a critical cell type that controls vascular permeability, promotes new vessel growth, regulates vascular tone, maintains the blood-retinal-barrier (**BRB**), and enables leukocyte extravasation[1-3]. Dysfunction of MVECs in the advanced stages of DR or AMD consequently leads to unregulated neovascularization and increased vascular permeability, causing hemorrhaging and inflammation that contribute towards rapid onset and severe visual impairment[4]. Current treatment options – photocoagulation of neovascular vessels and administration of anti-angiogenic agents –focus predominantly on slowing the growth of neovascular vessels in order to prevent hemorrhaging and vision loss. While these treatments are highly effective at slowing disease progression, they are destructive (e.g. photocoagulation) or have a high economic and social burden (e.g. anti-angiogenics), necessitating frequent invasive interventions throughout the patient’s life that are associated with increased risk of complications[5]. As such, the ability to modify MVECs via gene therapy in order to ameliorate dysfunction represents an alternative therapeutic option that could prevent the onset of sight-threatening complications if intervention is provided at an early stage. While numerous research groups have focused on identifying molecular targets for effective therapeutics in AMD and DR, the inability to efficiently mediate gene transfer to the affected MVECs has severely limited progress towards the development of a long-lasting gene therapy[6, 7].

Recombinant adeno-associated virus (r**AAV**) vectors have proven to be safe and effective tools for mediating gene transfer for the correction of inherited and acquired disorders in multiple organ systems, including the eye, liver and brain, in both pre-clinical animal models and human patients [8-12]. While multiple, naturally occurring serotypes exist that exhibit broad tissue tropism and the ability to transduce both dividing and non-dividing cells, alteration of the rAAV capsid through rational design or library screening has become a common approach to further alter cellular tropism, increase tissue penetrance, or provide the ability to evade cellular proteolytic degradation and host immunity [13-15].

Owing to their ubiquity throughout the body and involvement in multiple local and systemic diseases, MVECs have long provided an attractive, if not challenging target for genetic modification. Recently, a rational design approach towards enhancing the tropism of rAAV for MVECs involving the incorporation of endothelial cell (**EC**)-targeting peptides into surface-exposed sites of the capsid resulted in greater affinity of rAAV2/2-based mutant vectors for ECs isolated from the human vena cava and MVECs in the murine brain and lungs[16-21]. Similarly, recent studies have shown that incorporation of EC-targeting peptide insertions into the rAAV2/9 capsid can improve the transduction efficiency of human ECs harvested from the carotid artery *in vitro*[22, 23].

In this study, we follow a similar rational design approach towards vector engineering with the aim of enhancing the affinity of rAAV vectors for targeting MVECs within ocular tissues, including the choroid and retina. Specifically, we evaluate the transduction efficiencies of multiple rAAV2/2-, rAAV2/2[QuadYF+TV]-, and rAAV2/9-based vectors incorporating nine individual 7-mer EC-targeting peptides that have been demonstrated to significantly enhance tropism for MVECs in other tissues, including the mammalian brain and lung tissue[18, 19, 21, 24]. Following confirmation of packaging and transduction efficiency in *ex vivo* cultures of primary bovine MVECs, we assessed tropism of each EC-targeting (EC1-9 per serotype) capsid mutant vector and unmodified control expressing a ubiquitously expressing self-complementary green fluorescent protein (scGFP) reporter following intravitreal or intravenous injection in wild-type C57Bl/6J, using a combination of non-invasive ocular imaging and post-mortem histology.

## Methods

### Animals

Age-matched male and female C57BL/6 mice (N= 50, 25 males and 25 females) were purchased from the Jackson Laboratory (Bar Harbor, Maine). Mice were housed in the Biomedical Resource Center at the Medical College of Wisconsin (**MCW**) under a 12:12 light-dark cycle with food and water *ad libitum*. All animal protocols were reviewed and approved by the Medical College of Wisconsin Institutional Animal Care and Use Committee and conform with the National Institute of Health (**NIH**) and Association for Research in Vision and Ophthalmology (**ARVO**) guidelines for the Care and Use of Laboratory Animals.

### Isolation of Primary Bovine Retinal Endothelial Cells

Bovine eyes were harvested and purchased from Sierra-Medical (Whittier, CA, USA). A sterile work area was set up inside a laminar flow hood. Eyes were immersed in 1x betadine solution (Medline Industries, Northfield, IL) and rinsed in room temperature phosphate-buffered saline (**PBS**) (Gibco™, Dublin, Ireland) supplemented with 1% antibiotic/antimycotic solution. Eyes were dissected 5mm posterior to the limbus with a surgical blade. The retinae were gently removed from the eye cups, rinsed three times in PBS supplemented with 1% antibiotic/antimycotic solution, and collected in endothelial basal media (**EBM**) (Angio-Proteomie, Boston, MA) supplemented with 1% antibiotic/antimycotic solution. Retinae were homogenized in a rotary Teflon glass homogenizer, and the homogenate was filtered through a sterile 83µm mesh. Trapped tissue was collected and incubated in sterile EBM with collagenase (Gibco™, Dublin, Ireland) (0.5mg/mL), DNase I (Roche Diagnostic, Indianapolis, USA) (0.2mg/mL), and Pronase (Roche Diagnostic, Indianapolis, IN, USA) (0.2mg/mL) at 37 degrees Celsius with 5% CO2 for 45 minutes. After 45 minutes, the cell mixture was filtered through a 53µm mesh, and trapped tissue was collected in endothelial cell growth media (**EGM**) (Angio-Proteomie, Boston, MA, USA) supplemented with 1% antibiotic/antimycotic solution. Following centrifugation at 2000rpm for 10 minutes, the supernatant was discarded, and the cell pellet was resuspended in EGM. Cells were cultured on 0.1% gelatin coated plates at 37 degrees Celsius with 5% CO2.

### Recombinant rep-cap Plasmid Design

Mutant AAV2-, AAV2[QuadYF+TV]-, and AAV9-based rep-cap plasmids incorporating individual heptameric peptides (EC1-10) (pACG2-[EC1-10], pACG2-QuadYF+TV-[EC1-10], and pACG9-[EC1-10]) were generated through site-directed mutagenesis (NEB Q5 site-directed mutagenesis kit; New England Biolabs, Ipswich, MA, USA). Specifically, using individually designed forward and reverse primers (Tables 1, 2, and 3), the individual heptameric peptides were inserted between amino acids N587 and R588 of the AAV2 and AAV2[QuadYF+TV] *cap genes* and between amino acids A589 and Q590 of the AAV9 *cap* gene[22, 25].

**Table 1.**
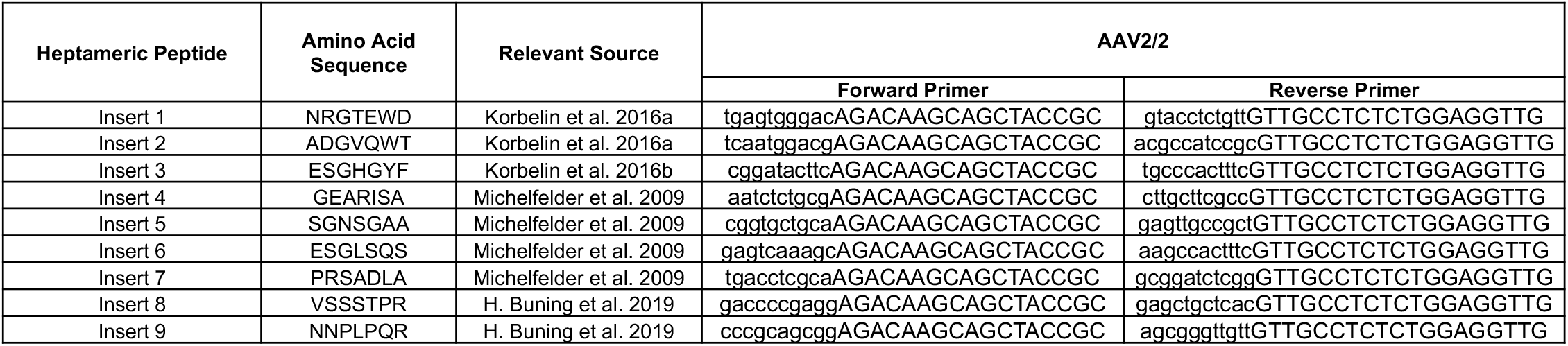
Primer Design for Site-Directed Mutagenesis of AAV2/2-based rep-cap Plasmids.

**Table 2.**
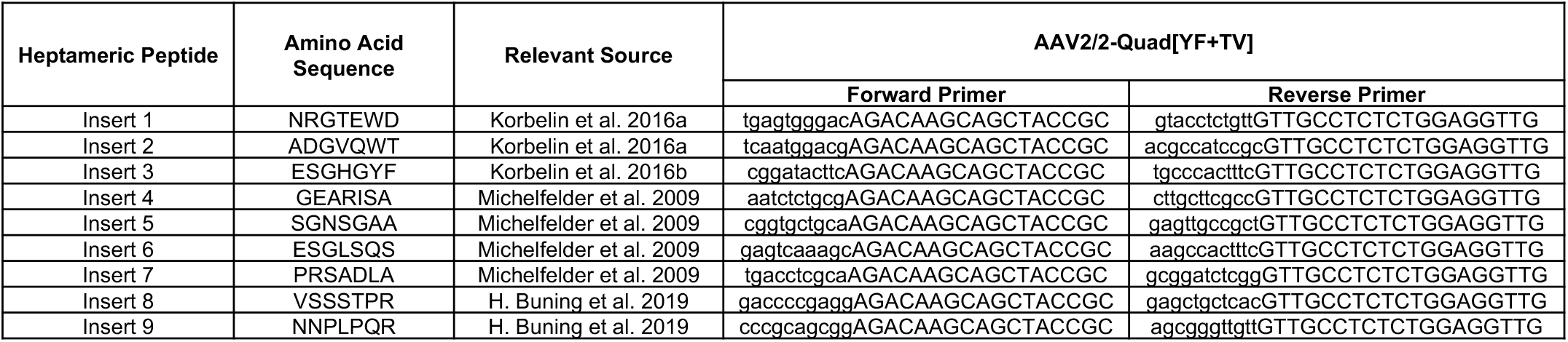
Primer Design for Site-Directed Mutagenesis of AAV2/2[QuadYF+TV]-based rep-cap Plasmids.

**Table 3.**
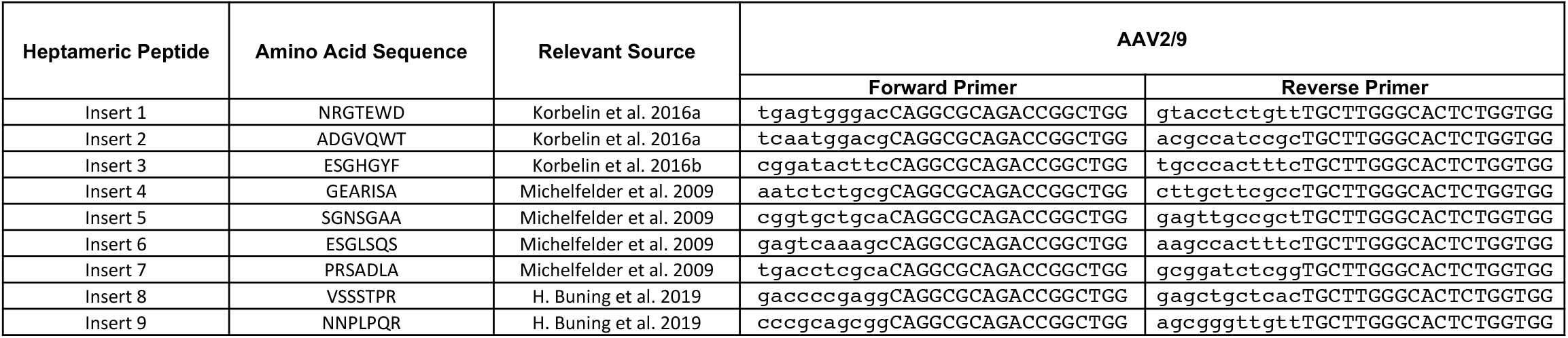
Primer Design for Site-Directed Mutagenesis of AAV2/9-based rep-cap Plasmids.

### In Vitro Screening of EC-Targeting Mutants

#### Vector Production

rAAV vector production was performed as previously described [26]. Briefly, HEK293T cells were triple transfected with an rAAV helper plasmid (pHelper); a ubiquitously expressing GFP reporter cassette (CBA-scGFP); and the appropriate EC-targeting mutant rep-cap plasmid (pACG2-[EC1-10], pACG2-QuadYF+TV-[EC1-10], or pACG9-[EC1-10]) or the unmodified rep-cap plasmids (pACG2, pACG2-QuadYF+TV, and pACG9) as control. Plasmids were transfected in equimolar ratios in high glucose DMEM (Gibco™, Dublin, Ireland) supplemented with 2% FBS and 1% antibiotic/antimycotic solution. HEK293T cells were harvested 72 hours post-transfection, and rAAV vector purification was performed by iodixanol density centrifugation and buffer exchange using 100kDa columns (Amicon, Darmstadt, Germany). The Virus was washed and eluted in Hank’s balanced salt solution (Gibco™, Dublin, Ireland) containing 0.014% Tween-20 (Sigma-Aldrich, St. Louis, MO, USA). Vector concentrations were determined using a PicoGreen assay (Thermo Fisher Scientific, Waltham, MA, USA).

### In Vitro Library Screen and Flow Cytometry

Primary bovine retinal microvascular endothelial cells (**BMVECs**) were cultured at passage number two (P=2) in 0.1% gelatin-coated 24-well plates with EGM at 37 degrees Celsius with 5% CO_2_. At 70% confluency, BVECs were infected in triplicates (75,000 MOI) with the mutant rAAV2/2-[EC1-10].CBA-sc.GFP, rAAV2/2[QuadYF+TV]-[EC1-10].CBA-sc.GFP, or rAAV2/9-[EC1-10].CBA-sc.GFP vectors, or unmodified rAAV2/2.CBA-sc.GFP, rAAV2/2[QuadYF+TV].CBA-sc.GFP, or rAAV2/9.CBA-sc.GFP vectors as control.

After 96 hours, transduced BMVEC cells were dissociated with TrypLE Express (Gibco™, Dublin, Ireland) into single cells and washed with PBS containing 10% fetal bovine serum (**FBS**) (Gibco™, Dublin, Ireland) and centrifuged at 200xg for 5 minutes. The pelleted cells were blocked with 0.5 ml incubation buffer (0.5% BSA, PBS) for 15 minutes at room temperature in a 5 mL round bottom polystyrene tube (Corning, Tewksbury, MA, USA). Next, Cells were rinsed in PBS containing 10% FBS and centrifuged at 200xg for 5 minutes prior to labeling them with the anti-mouse CD31 monoclonal primary antibody (5ug/1×10^6^ cells, Invitrogen, Carlsbad, CA, USA) in incubation buffer for 90 minutes at room temperature (**RT**). After incubation, cells were rinsed (3 times) with 2 ml incubation buffer and resuspended with anti-mouse Alexa Fluor 647 secondary antibody (1:2000, Invitrogen, Carlsbad, CA, USA) for 60 minutes at RT. The cells were then rinsed in the incubation buffer (3 times) and resuspended in the Flowcytometry buffer (Ca/Mg2+ Free PBS, 2% FBS, 0.1% Sodium Azide). Cells were quantified by LSR-II (BD Biosciences, San Jose, CA, USA), and, using flow cytometry analysis software (FlowJo v10.7.2), transduction efficiency was quantified as the percentage of GFP+CD31+ cells normalized to the total percentage of CD31+ cells in the sample.

### *In Vivo* Assessment of Transduction and Tissue Tropism

#### Virus Production

High titer rAAV2/2.CBA-eGFP, rAAV2/9.CBA-eGFP, rAAV2/2[QuadYF+TV].CBA-eGFP. rAAV2/2-EC5.CBA-eGFP, rAAV2/9-EC5.CBA-eGFP, rAAV2/2[QuadYF+TV]-EC.CBA-eGFP.were manufactured by Dr. Boye at the University of Florida College of Medicine and eluted in TMN200-P (200mM NaCl, 1mM MgCl2, 20mM Tris, pH adjusted to 8.0 and supplemented with 0.001% F68 Pluronic).

#### Intravitreal and Retro-orbital Injections

Intravitreal and retro-orbital injections were performed under isoflurane anesthesia. Anesthesia was induced using 5% isoflurane in 100% oxygen before being reduced to 2% isoflurane for maintenance. Pupils were dilated using a combination of 2.5% phenylephrine (Paragon BioTeck, Portland, OR) and 1% tropicamide (Akorn, Lake Forest, IL). The moisture of the eye was maintained during imaging using Systane Ultra lubricant eye drops (Alcon Inc., Fort Worth, TX).

Using a 33-gauge needle and Hamilton syringe, cohorts of age-matched male and female C57BL/6 mice (N=6 mice for each vector, 3 male and 3 female) received 2μL of any one rAAV vector (∼2.5×10^12^vg/mL) through bilateral intravitreal injections. Using a zero-volume syringe and 28-gauge needle (BD PrecisionGlide Needle, BD, Franklin Lakes, NJ, USA), age-matched C57BL/6 mice (N=2, one male and one female) received 150μL of any one rAAV vector (∼2.5×10^12^ vg/mL) through retro-orbital injection.

#### Ocular imaging

Four months post-injection, all study animals were imaged using a custom multiline confocal scanning laser ophthalmoscope (cSLO; modified Spectralis HRA, Heidelberg Engineering, Heidelberg, Germany). Camera alignment was performed using near-infrared (820nm) reflectance imaging. Retinal images of vector-derived GFP fluorescence (482nm excitation; 502-537nm bandpass) were obtained and processed for false color using ImageJ software (**NIH**).

#### Cryosection and immunostaining

Four weeks postinjection, mice were euthanized by transcardial perfusion under terminal anesthesia. In short, anesthetized mice were fixed in a supine position, and a thoracotomy was performed. First, 10 ml of PBS followed by 10 ml of 4% PFA was perfused into the left ventricle, and the right atrium was resected. As the fixative (PFA) enters, the body stiffens, and the liver turns pale, indicating a successful perfusion procedure. After perfusion, organs were harvested, cryoprotected in 30% sucrose overnight at 4°C, embedded in the optimal cutting temperature medium, and stored at -80°C until cryo-sectioned. Cryo-sections (12 mm) were made with a cryostat microtome system (Leica, Germany) and stored at -20° C before antibody staining. Sections were permeabilized with 1x PBS containing 0.2% Triton-X for 20 minutes, stained with nuclear stain DAPI (1 in 10,000), and washed with 1xPBS (3 times) for 15 mins. Slides were imaged for native GFP signal and DAPI nuclear stain on a fluorescence microscope (EVOS M5000, Invitrogen).

## Results

### Screening transduction efficiency of rAAV vectors containing EC-targeting heptameric peptide insertions in primary bovine VECs

Insertion of heptameric peptides between amino acid positions 587/588 in rAAV2/2-based capsids interferes with two (R585 and R588) of the five positively charged amino acids that has been identified for HSPG binding [27, 28]. However, a net positive charge of the inserted peptide, i.e., “bulkiness” of the inserted residues, and proximity of an arginine or alternative positively-charged residue at the R588 position may reconstitute HSPG binding [29]. Heptameric peptide inserts EC1, EC2, EC3, and EC6 have a net negative charge at pH 7, while inserts EC4, EC5, and EC7 have a net neutral charge, and EC8 and EC9 have net positive charges (Table 4). Inserts EC8 and EC9 also have a positively charged arginine residue within one amino acid position of R588, while EC3 carries a positively charged histidine residue separated by three amino acid residues from R588. As a result of differences in charge and ‘bulkiness’ of each 7-mer peptide, we anticipated that their incorporation into rAAV2/2, rAAV2/2[QuadYF-TV] or rAAV2/9 [at positions 588 (2/2 and 2/2[QuadYF-TV] or 589 (2/9) of the virus protein (VP1-3)] virions may adversely affect packaging and infectivity of each EC mutant (EC1 – EC9) compared to unmodified controls. Following optimization of multiplicity of infection (MOI) in HEK293T (MOI=50,000) and primary bovine MVECs (BMVECs; MOI=75,000) for each serotype (rAAV2/2, rAAV2/2[QuadYF-TV] and rAAV2/9), we confirmed that the preparations of each EC mutant vector (EC1-9) were infectious (Figs 1A and B), as evidenced by robust GFP expression after a period of 72-hours (Supplemental Fig 1). Subsequently, primary BMVECs were transduced with rAAV2/2-EC1-9, rAAV2/2[QuadYF-TV]-EC1-9 or rAAV2/9-EC1-9 vectors (N=3 wells per mutant, N=27 mutants) or unmodified rAAV2/2, rAAV2/2[QuadYF-TV] or rAAV2/9 (N=3 wells per capsid; Fig 2). Transduction efficiency was quantified 96 hours after vector supplementation by calculating the percentage of GFP^+^CD31^+^ cells normalized to the total percentage of CD31^+^ cells in each sample, allowing the fold-change decrease/increase in transduction efficiency of each mutant vector to be compared to the transduction efficiency of its respective, unmodified parental serotype. rAAV2/2-based mutants with fold-changes greater than 1x included: rAAV2/2-EC1 (2.1x), rAAV2/2-EC5 (2.72x), rAAV2/2-EC6 (1.57x), (Fig 3E). Notable rAAV2/2[QuadYF+TV]-based mutants included: rAAV2/2[QuadYF+TV]-EC2 (1.15x), rAAV2/2[QuadYF+TV]-EC4 (2.77x), rAAV2/2[QuadYF+TV]-EC5 (3.68x), rAAV2/2[QuadYF+TV]-EC6 (7.82x) (Fig 3E). Notable rAAV2/9-based mutants included: rAAV2/9-EC5 (1.72) (Fig 3E).

**Table 4.** Net-Charge of EC-Targeting Heptameric Peptides.

| Heptameric Insert | Amino Acid Sequence | Net-Charge<br>At pH 7 |
| --- | --- | --- |
| EC1 | NRGTEWD | -1 |
| EC2 | ADGVQWT | -1 |
| EC3 | ESGHGYF | -0.9 |
| EC4 | GEARISA | 0 |
| EC5 | SGNSGAA | 0 |
| EC6 | ESGLSQS | -1 |
| EC7 | PRSADLA | 0 |
| EC8 | VSSSTPR | 1 |
| EC9 | NNPLPQR | 1 |

**Figure 1.**
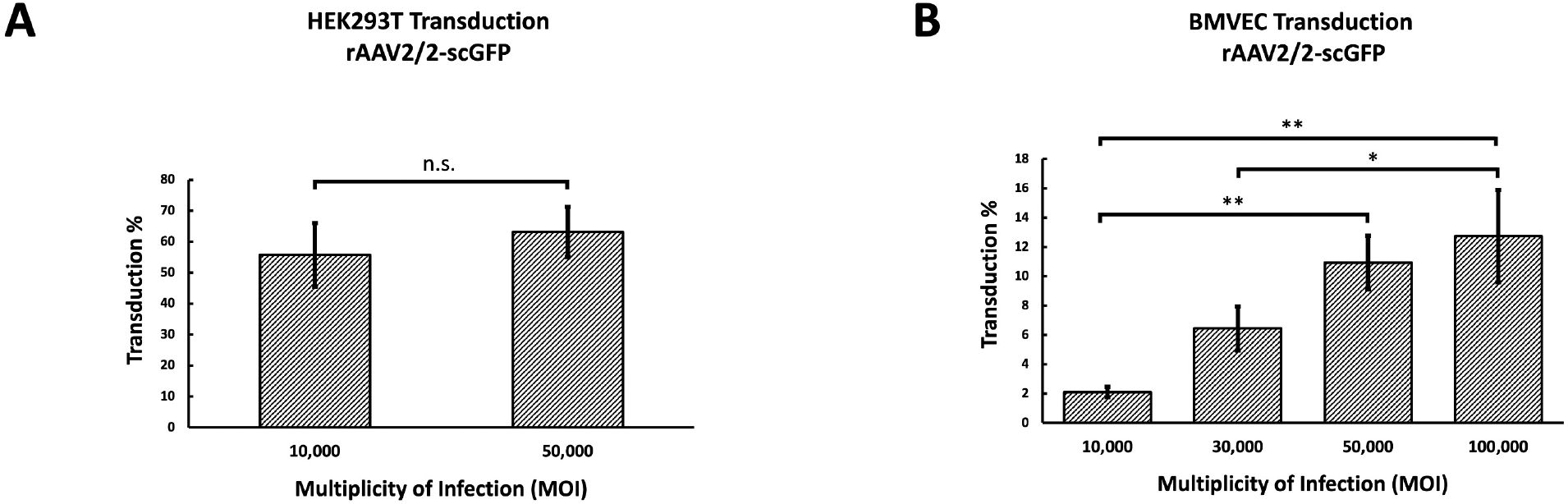
Optimizing MOI and Validating Infectivity. **(A)** HEK293T cells were seeded in 24-well plates (∼500,000 cells/well), and transduced with rAAV2/2.CBA-scGFP at 10,000 MOI (N=3 wells) and 50,000 MOI (N=3). Percent transduction was quantified by flow cytometry as the percentage of GFP+ cells. A paired t-test revealed that the difference in percent transduction between 10,000 MOI and 50,000 MOI was not significant (n.s.) (B). Primary BMVECs were seeded in 24-well plates (∼100,000 cells/well), and cells were transduced with rAAV2/2.CBA-scGFP at 10,000, 30,000, 50,000, and 100,000 MOI (N=3 wells per MOI). One-way ANOVA with post-hoc Tukey test revealed significant dose-dependent differences in the percent transduction of BMVECs (*p<0.05, **p<0.01). Based on the findings in Panel A, the infectivity of rAAV2/2, rAAV2/2[QuadYF+TV], rAAV2/9, and respective mutant vectors packaging a ubiquitously expressing GFP reporter cassette were assessed upon transducing HEK293T cells at 10,000 MOI

**Figure 2.**
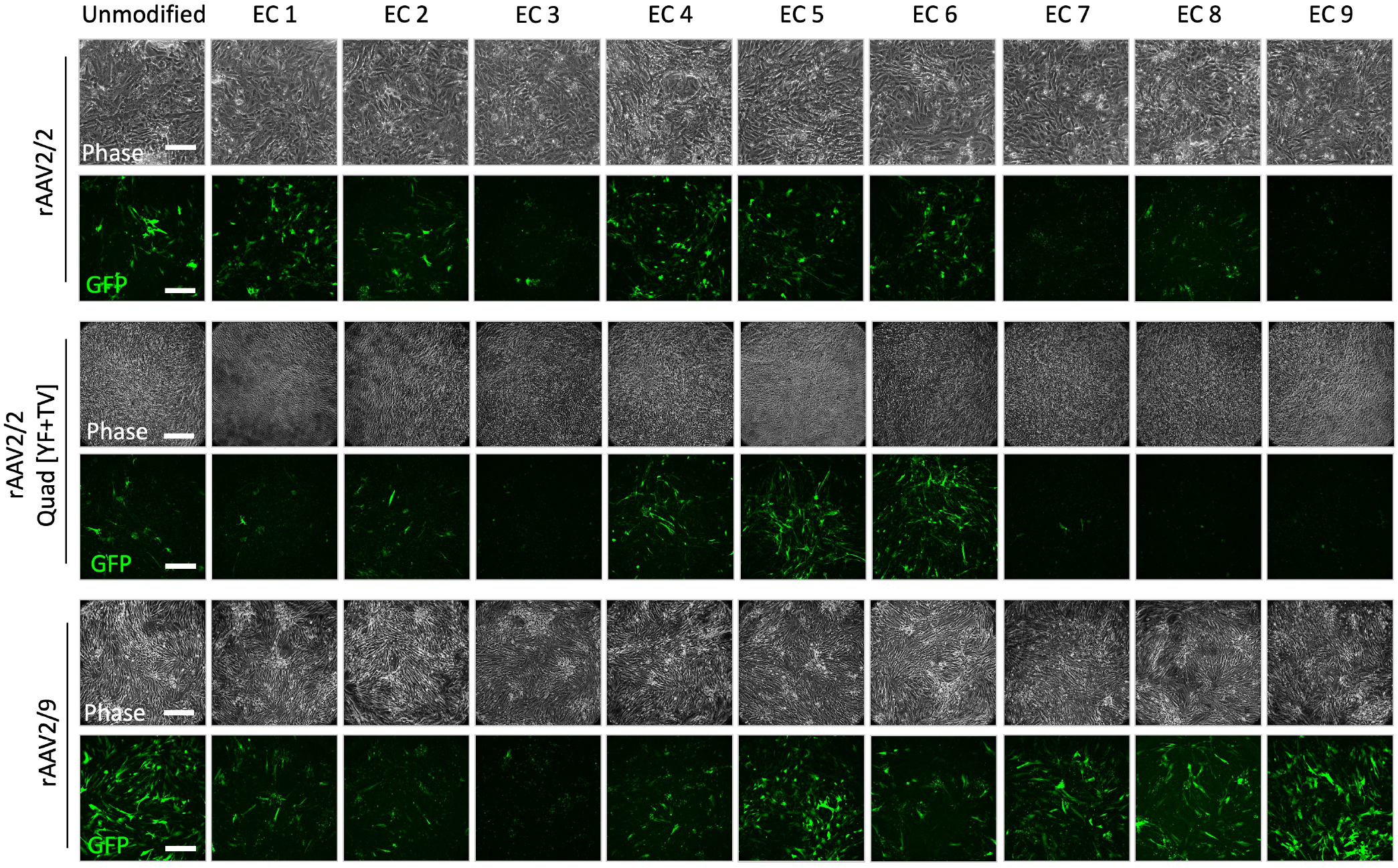
In Vitro Screening of rAAV2/2, rAAV2/2[QuadYF+TV] and rAAV2/9-based EC Mutants. Based on the findings in Figure 1B, primary BMVECs were infected in triplicates with the vectors at 75,000 MOI. Scale bar: 100μm.

**Figure 3.**
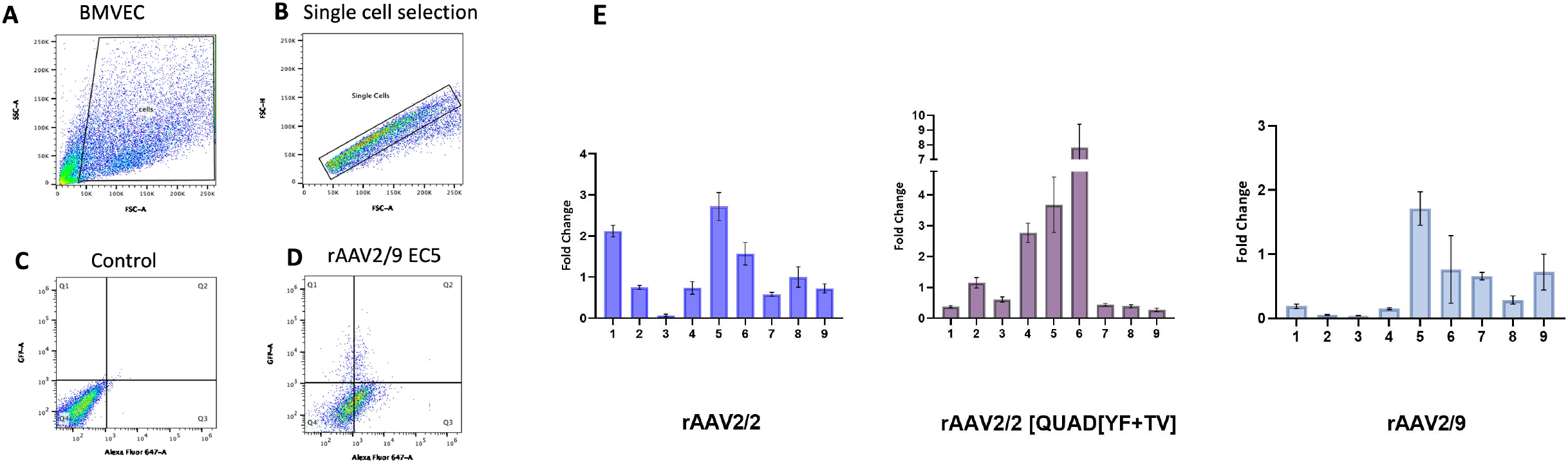
Transduction efficiency by Flowcytometry. (A) Representative Flow Cytometry Gating for BMVEC stained with CD31 Antibody. Flow cytometry was performed following rAAV transduction. BMVEC were analyzed after debris exclusion (A) and single-cell selection (B). Untransduced BMVEC cells were used as the gating control (C), Q1 = GFP+ve cells only, Q2= GFP+ve and CD31+ve cells, Q3=CD31+ve cells only, and Q4=GFP-ve and CD31-ve cells (D). (E) Transduction efficiency was quantified as the total percentage of GFP+CD31+ cells normalized to the percentage of CD31+ cells in each well. The transduction efficiency of each mutant EC-targeting vector was compared to unmodified rAAV2/2, rAAV2/2[QuadYF+TV], and rAAV2/9 as fold-change (n=3).

Interestingly, rAAV vectors incorporating EC5 performed better than their respective unmodified control vectors for all serotypes evaluated, and therefore, vectors incorporating EC5 were selected for further studies to determine their transduction and tissue tropism, *in vivo*.

### Retinal transduction following retro-orbital or intravitreal injection of EC-mutant vectors

After determining that EC5-containing capsid mutant vectors out-performed the parental strains for all serotypes, rAAV2/2-EC5, rAAV2/2[QuadYF-TV]-EC5 and rAAV2/9-EC5 mutants and unmodified control vectors were administered systemically to characterize both the ocular and systemic tropism. Unilateral retro-orbital injections (100μL, ∼3.75×10^11^vg per mouse) were performed in age-matched C57BL/6 mice (N = 6 male, N=6 female) with each vector packaging a ubiquitously expressing enhanced GFP (**eGFP**) reporter cassette. Twelve-weeks post-injection, eyes were imaged using confocal scanning laser ophthalmoscopy (cSLO) with near-infrared reflectance (NIR; 820nm) and autofluorescence (AF; 488nm) modalities to assess for evidence of ocular damage or vector-mediated eGFP expression, respectively (Fig 4). NIR imaging (Supplemental Fig 2) demonstrated no evidence of ocular injury following high volume vector injection into the retro-orbital venous sinus. AF imaging revealed widespread punctate signals throughout the neural retina in both unmodified and EC mutant injected eyes, potentially indicative of macrophage infiltration. In rAAV2/2-EC5, rAAV2/2[QuadYF-TV]-EC5 and rAAV2/9-EC5 mutant injected animals, increased fluorescence signal was observed along the vessel walls (red arrows, Fig 4), specifically in the vicinity of the optic nerve head, indicating moderately increased levels of retinal vascular transduction compared to unmodified rAAV2/2, rAAV2/2[QuadYF-TV] and rAAV2/9 injected animals.

**Figure 4.**
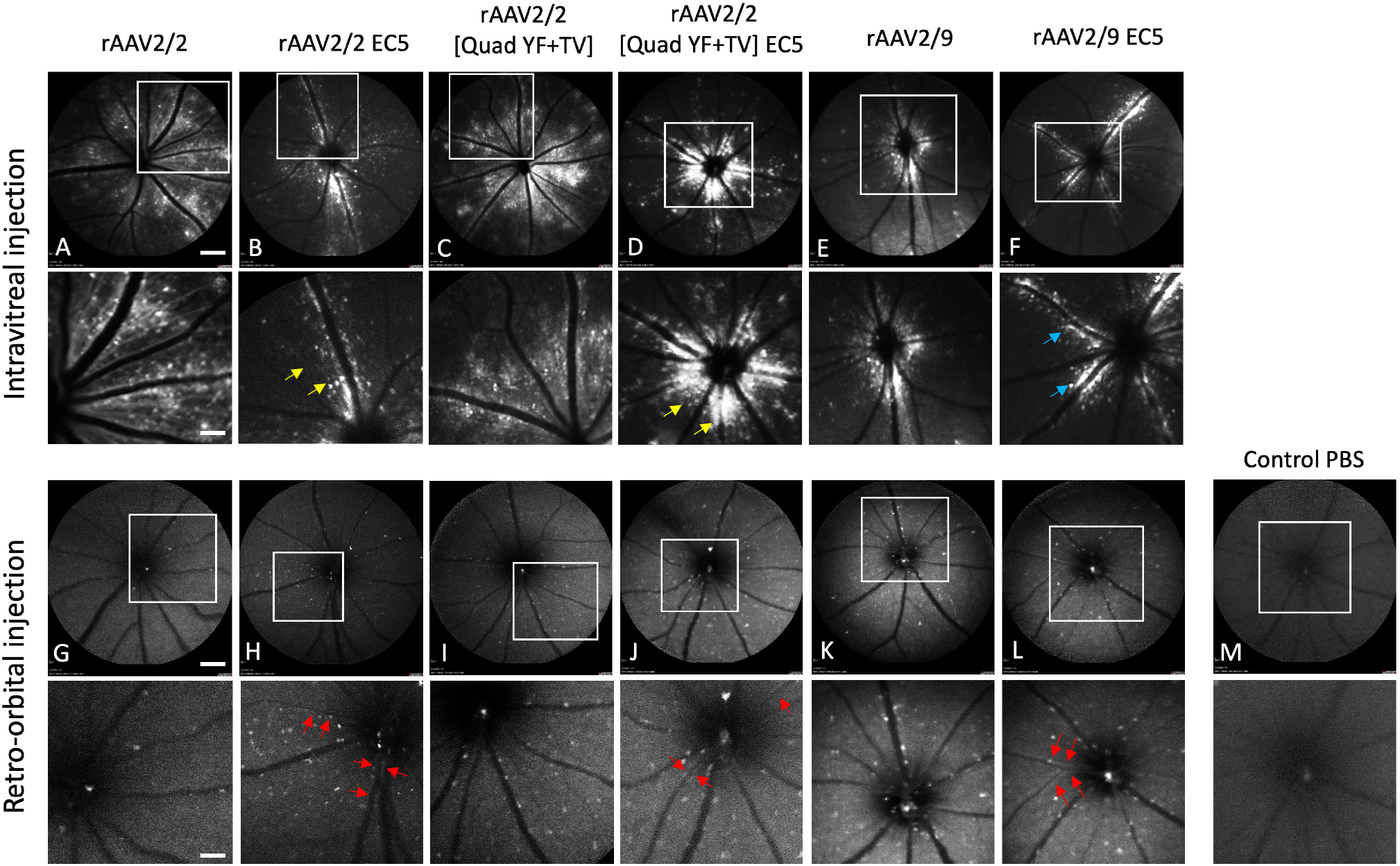
In vivo assessment of rAAV-mediated GFP Expression in mice retina. C57BL/6 mice received 2ul of mutant vectors containing EC5 and the unmodified vector as control through bilateral intravitreal injections (A-F) and unilateral retro-orbital injections (G-L), and PBS control (M). GFP expression were imaged four months post-injection using a multi-line cSLO. Note the GFP expression (top row) with the indicated insets magnified (bottom row) Red arrows indicate retinal vessel (vascular) transduction, Yellow arrows indicate the transduction of cells closer to the optic discs, and Blue arrows indicate GFP expression from the cells closer to the major retinal blood vessels. Scale bar: 800μm.

Intravitreally injected eyes revealed differential expression patterns in all EC-mutant injected eyes relative to unmodified controls. Specifically, rAAV2/2-EC5 and rAAV2/2[QuadYF-TV]-EC5 injected eyes demonstrated reduced transduction of retinal ganglion cells, as identified by fewer axonal projections extending from the periphery towards the optic disc (yellow arrows, Fig 4), and more focal transduction centered around the optic nerve head than in eyes injected with unmodified vectors (Fig 4). rAAV2/9-EC5 mutant injected eyes demonstrated the most altered transduction, with the greatest concentration of GFP signal occurring in cells immediately surrounding the major retinal blood vessels (blue arrows, Fig 4).

### Systemic transduction following retro-orbital or intravitreal injection of EC-mutant vectors

In addition to determining the extent of ocular vascular transduction following retro-orbiral or intravitreal injection of rAAV2/2-EC5, rAAV2/2[QuadYF-TV]-EC5 or rAAV2/9-EC5, several organs (notably, the liver, heart and lungs) were harvested from all animals and examined for evidence of GFP expression localized in blood vessels or proximal structures. GFP expression was observed in the liver in all retro-orbital injected animals, but was especially prominent in those administered rAAV2/2[QuadYF-TV]-EC5 or rAAV2/9-EC5, with high levels of expression surrounding the hepatic vessels (white arrows) and bile ducts (Fig 5, bottom panel, J & L). Interestingly, rAAV2/9-EC5 animals injected intravitreally also demonstrated evidence of GFP expression in the liver, indicating widespread virion escape from the eye into the systemic circulation (Fig 5, upper panel, F & L). In addition to targeting the liver, rAAV2/2[QuadYF-TV]-EC5 and rAAV2/9-EC5 mutants demonstrated a moderate ability to transduce lung tissue following retro-orbital injection (Fig 6, bottom panel, J and K). Meanwhile rAAV2/2[QuadYF-TV]-EC5 showed dramatically increased tropism for cardiac muscle compared to the unmodified rAAV2/2[QuadYF-TV] parent serotype (Fig 7, bottom panel, D and J). Intravitreal injection of all vectors resulted in an absence of observable fluorescence signal in the lungs and hearts of all animals.

**Figure 5.**
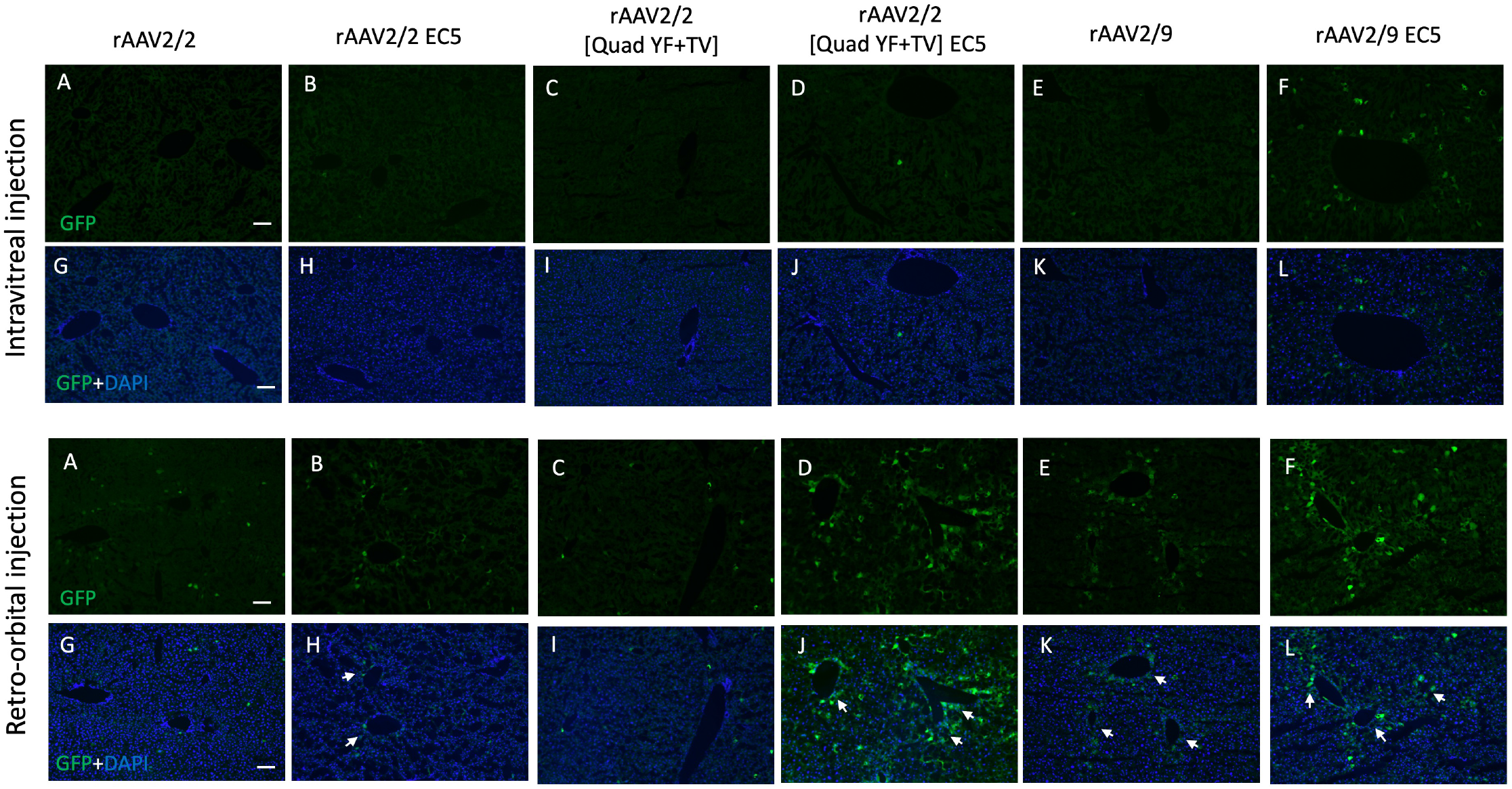
rAAV-mediated transduction in murine liver by Intravitreal and Retro-orbital delivery. C57BL/6 mice received 2ul of mutant vectors containing EC5 and the unmodified vector as control through bilateral intravitreal injections (Top Panel) and unilateral retro-orbital injections (Bottom Panel). After four months post-injection, Immunofluorescence of cryosectioned liver sections was imaged for GFP signal (A-F) with counterstaining of DAPI (G-L). Note: White arrows indicate cells having high GFP expression surrounding the hepatic and bile ducts.

**Figure 6.**
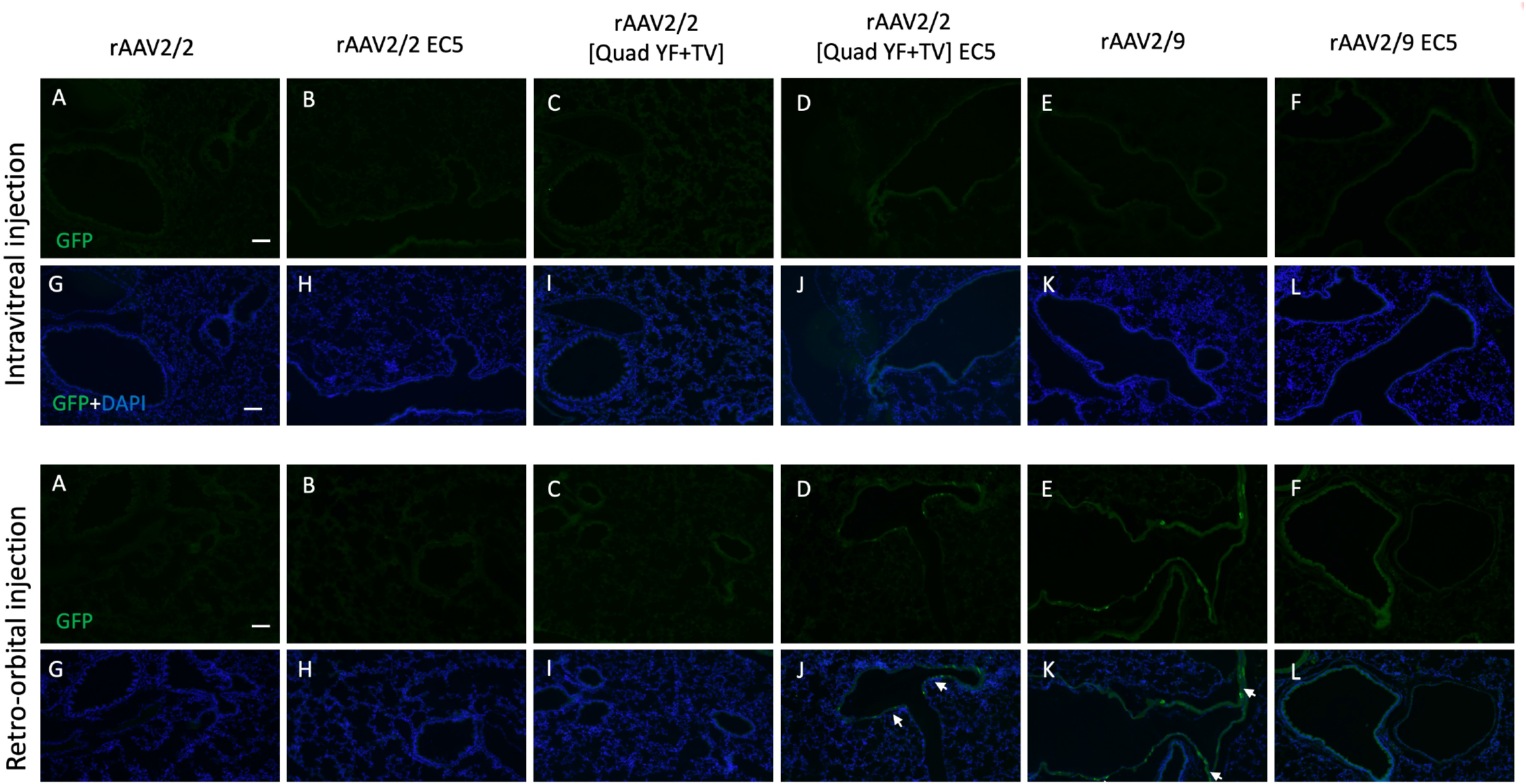
rAAV-mediated transduction in murine lung by Intravitreal and Retro-orbital delivery. C57BL/6 mice received 2ul of mutant vectors containing EC5 and the unmodified vector as control through bilateral intravitreal injections (Top Panel) and unilateral retro-orbital injections (Bottom Panel). After four months post-injection, Immunofluorescence of cryosectioned lung sections was imaged for GFP signal (A-F) with counterstaining of DAPI (G-L). Note: White arrows indicate cells having GFP expression surrounding the blood vessels.

**Figure 7.**
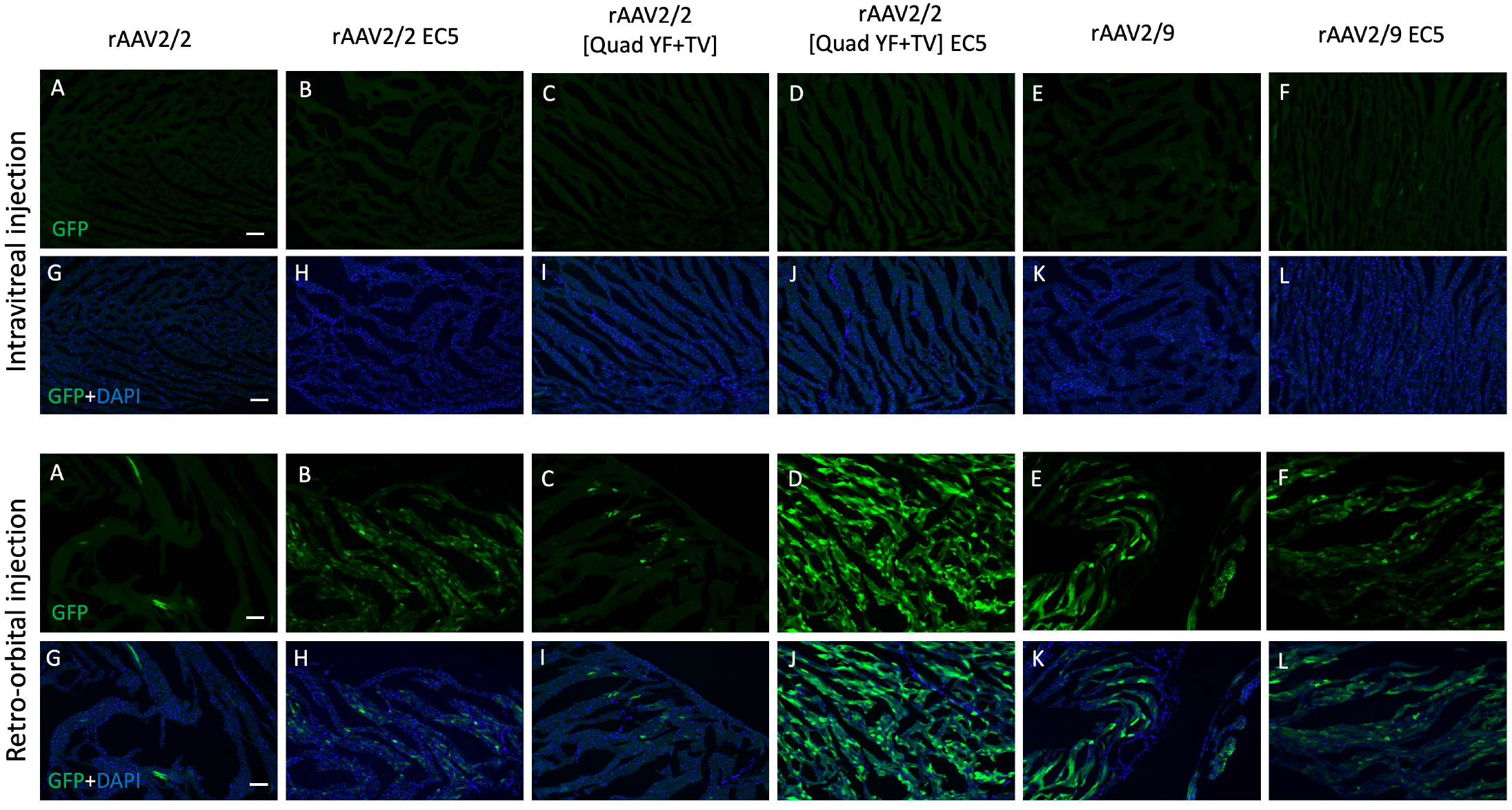
rAAV-mediated transduction in murine heart by Intravitreal and Retro-orbital delivery. C57BL/6 mice received 2ul of mutant vectors containing EC5 and the unmodified vector as control through bilateral intravitreal injections (Top Panel) and unilateral retro-orbital injections (Bottom Panel). After four months post-injection, Immunofluorescence of cryosectioned heart sections was imaged for GFP signal (A-F) with counterstaining of DAPI (G-L).

## Discussion

rAAV vectors have successfully enabled safe and efficient gene delivery for the treatment of various ocular diseases, such as Leber’s congenital amaurosis (**LCA**)[9, 30-32], Leber’s hereditary optic neuropathy[33], and choroideremia[34]. Despite progress, the application of rAAV vectors for the treatment of complex retinal vascular diseases is severely limited by the absence of rAAV vectors capable of robustly transducing MVECs, a critical cell type, the dysfunction of which underlies pathology in several diseases, including diabetic retinopathy and age-related macular degeneration. Herein, we screened multiple capsid mutant rAAV vectors containing heptameric peptide insertions designed to increase endothelial cell targeting and uptake. Initial evaluations were performed *in vitro* using HEK293T and BMVECs cultures to determine infectivity and identify mutants with the greatest tropism for CD31+ endothelial cells, and subsequently *in vivo* following either intravitreal or retro-orbital (intravenous) injection in wild-type mice using a combination of non-invasive imaging and post-mortem histology. Engraftment of the EC5 peptide (SGNSGAA) into the capsid of rAAV2/2, rAAV2/2[QuadYF-TV] and rAAV2/9 resulted in infectious virions with altered tropism, including moderately increased transduction of the retinal vasculature (rAAV2/2[QuadYF-TV]-EC5) and liver vasculature (rAAV2/2[QuadYF-TV]-EC5 and rAAV2/9-EC5), and dramatic increased transduction of heart muscle (rAAV2/2[QuadYF-TV]-EC5) following intravenous administration.

Insertion of the heptameric peptides was carried out between amino acid positions N587 and R588 of the rAAV2/2 and rAAV2/2[QuadYF-TV] *cap* genes (A589 and Q590 in rAAV2/9), which is known to cause disruption of the AAV2 virion’s HSPG-binding motif and impair infection; however, the combination of reduced HSPG-binding affinity and insertion of a cell-targeting peptide is likely to yield a vector with re-directed tropism compared to the parental strain[35]. Indeed, several studies have previously identified rAAV capsid mutant vectors demonstrating enhanced transduction of MVECs in mammalian lungs and the brain following the insertion of similar EC-targeting heptameric peptides into the surface-exposed loop domains of the virion’s structural proteins; however, none of these studies have described the transduction efficiencies of such EC-targeting rAAV mutant vectors in retinal MVECs [16-21].

The heptameric peptides used for this study were selected based on their ability to transduce MVECs and macrovascular ECs with high efficiency in non-ocular systems. For example, Körbelin et al. (2016) demonstrated systemic administration of rAAV2-based vectors incorporating heptameric peptides with sequence motifs XXGXXWX substantially improved transduction of murine brain microvasculature (peptide inserts EC1 and EC2) and cardiac tissue (EC2)[21]. Körbelin et al. (2016b) also identified a rAAV2-based vector incorporating EC3 that mediated strong lung MVEC-specific expression of GFP (>200 times greater than rAAV2/2) following systemic administration in mice[21]. Interestingly however, in the present study, vectors incorporating this same peptide insertion (rAAV2/2-EC3, rAAV2/9-EC3, and rAAV2/2[QuadYF+TV]-EC3) were amongst the least efficient at targeting primary BMVECs *in vitro* (Figs 2 and 3), with all mutants demonstrating lower transduction than their unmodified parental strains regardless of serotype. While we have not evaluated the rAAV EC-mutant vectors on MVECs harvested from organs other than the retina – as our primary focus was to increase gene delivery efficiency to the retinal vasculature – this discrepancy highlights the heterogeneity of endothelial cells throughout the body and the importance of screening on the relevant primary tissues.

Based on the *in vitro* screening there appears to be a broad discrepancy between the GFP expression observed on fluorescent imaging (Fig 2) and flow cytometry data (Fig 3), where for example rAAV2/2[QuadYF-TV]-EC6 does not visibly appear to exhibit significantly greater levels of transduction than either rAAV2/2-EC6 and rAAV2/9-EC6 mutants, but demonstrates a ∼7.5-fold increase in BMVEC transduction when measured via flow cytometry. This discrepancy almost certainly arises from the primary BMVECs cultures being comprised of a mixed population of ECs and non-ECs arising from the size exclusion approach used for isolation following dissociation of the retinal vasculature ([36, 37]). As such, GFP expression observed via fluorescence microscopy may arise either from transduction of BMVECs or contaminating neuronal/glial cells. By contrast, flow cytometry was performed using co-staining with the CD31 endothelial-specific cell-marker and as such, we believe that this data more accurately represents the affinity of each EC mutant for retinal endothelial cells within the retina.

Based on the performance of EC-targeting peptides at transducing primary cells isolated from bovine retinae *in vitro*, we identified insert EC5 as the most promising EC-targeting peptide for subsequent *in vivo* assessment. Notably, while the EC6 peptide insertion (ESGLSQS) demonstrated up ∼7.5-fold increase in BMVECs when engrafted onto the rAAV2/2[QuadYF-TV] capsid, only the EC5 peptide insertion conferred increased transduction efficiency across all three serotypes examined, indicating it may mediate a broad serotype-independent increase in endothelial cell affinity. Following intravitreal injection of EC mutants into wild-type C57BL/6j mice, we observed a subtle alteration of tropism on cSLO imaging, with fluorescence signal more concentrated around the larger blood vessels and optic disc; however, no evidence of vascular or endothelial transduction was noted for any serotype. This finding was largely expected, where inclusion of any EC-targeting peptide into the capsid is likely only to increase transduction when the virion has direct access the endothelial cell membrane from the lumen (i.e., from within the lumen) and is not likely to increase vascular penetrance from the extra-luminal vitreous aspect.

By contrast, intravenous injection of EC-mutants resulted in some evidence of vascular transduction on cSLO imaging, with increased fluorescence signal apparent in the vessel walls localized proximal to the optic disc, where virions would be expected to enter the eye at the highest dose following injection into the retro-orbital venous sinus.

The most dramatic increase in transduction efficiency was observed not in the eye, but in the heart of rAAV2/2[QuadYF-TV]-EC5 injected animals, with robust and widespread GFP expression throughout the cardiac muscle compared to the parental rAAV2/2[QuadYF-TV] vector. The EC5 mutant was initially identified during a biopanning experiment aimed at targeting solid tumors in breast cancer, alongside two other peptide insertions also evaluated herein, namely EC4 (GEARISA) and EC6 (ESGLSQS).[18] While insertions EC4 and EC6 were observed in Michelfelder et al (2009) to result in increased heart transduction, mutant EC5 (SGNSGAA) was not evaluated beyond the initial library screen, and as such the increased heart muscle transduction observed herein with this peptide insertion was not identified. That dramatically increased heart transduction was observed following EC5 insertion into rAV2/2[QuadYF-TV], but not unmodified rAAV2/2, indicating that further disruption of HSPG-binding, which is already weakened in the former, may be necessary for cardiac tropism. Indeed, a previous study has shown that enhanced heart transduction was observed upon modification of the capsid VP3 region at position R588 to reduce HSPG binding and this region contributes to heart tropism [38] [16, 39]. However, that Michelfelder et al (2009) noted no significant upregulation of cardiac tropism when a random peptide (VRRPRFW) was inserted at position R588 of rAAV2/2 strongly suggests that the hydrophobicity (EC5 = neutral; random = hydrophobic) or overall net-charge (EC5 = 0.00; random = 3.00, both at pH7.0) of the insert is also important for increasing cardiac tropism, rather than simply disrupting HSPG binding affinity (Table 4).

## ACKNOWLEDGMENTS

The authors thank Christine Skumatz, Lisa King, Joseph Thulin, and the Biomedical Resource Center staff for the animal care. Grants from the National Eye Institute supported this work……

## AUTHOR CONTRIBUTIONS

R.P, D.P, and D.M.L. designed and conducted the experiments and wrote the paper. S.E.B. and S.L.B provided purified vectors for *in vivo* injections and reviewed the manuscript.

## Figure and Table legends

**Supplemental Figure 1.**
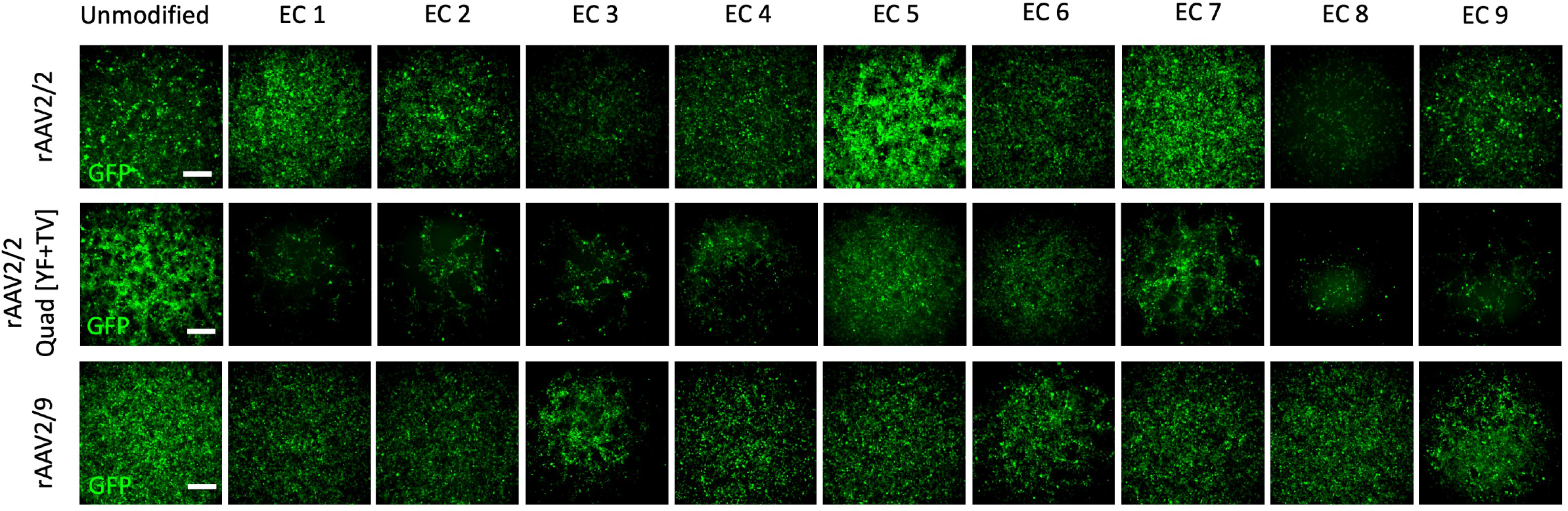
Screening of transduction efficiency in BMVEC. All vectors at 75,000 MOI were introduced to BMVEC and observed under a Fluorescence microscope for GFP expression after 72 hours. All the control vectors could transduce the BMVEC (GFP positive); however, GFP expression varied among the mutant vectors. Scale bar: 150μm.

**Supplemental Figure 2.**
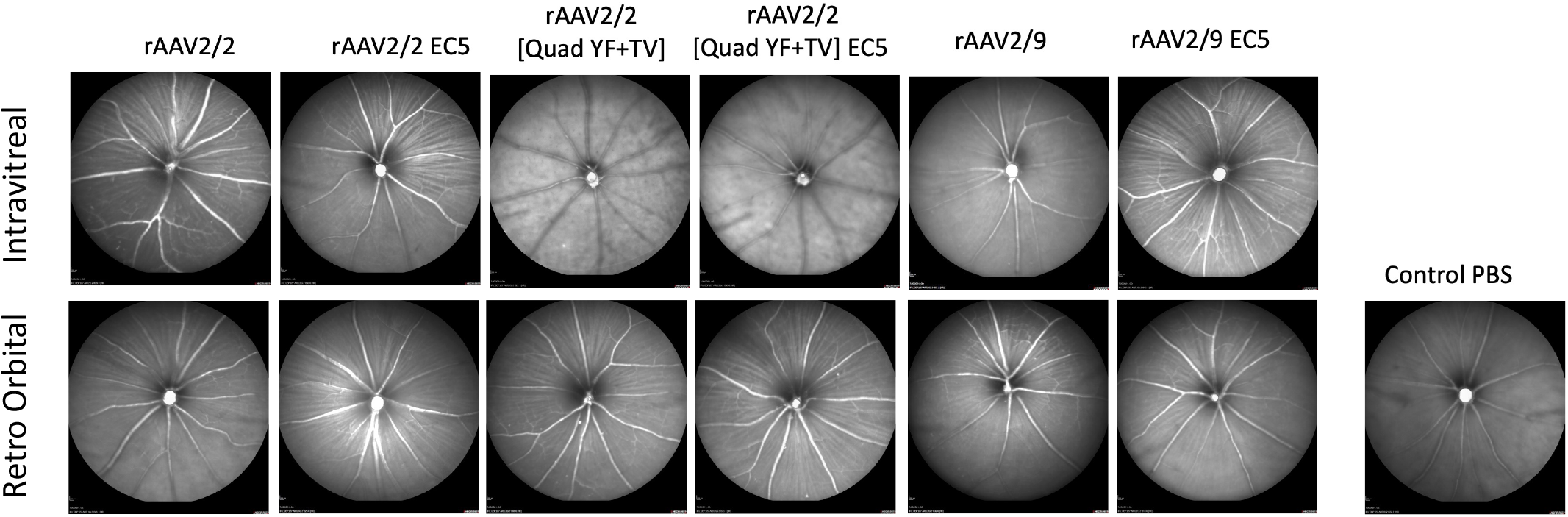
In vivo assessment of rAAV injections in mice retina. Near-infrared (820nm) reflectance imaging was performed using Confocal scanning laser ophthalmoscopy (cSLO) four months post-injection, and No evidence of ocular damage was observed. The above figure represents the cSLO images of mice eyes that received Intravitreal and Retro Orbital injections (2ul each) of all vectors and control (PBS).

